# Structure supports function: informing directed and dynamic functional connectivity with anatomical priors

**DOI:** 10.1101/2021.05.11.443529

**Authors:** David Pascucci, Maria Rubega, Joan Rué-Queralt, Sebastien Tourbier, Patric Hagmann, Gijs Plomp

## Abstract

The dynamic repertoire of functional brain networks is constrained by the underlying topology of structural connections: the lack of a direct structural link between two brain regions prevents direct functional interactions. Despite the intrinsic relationship between structural (SC) and functional connectivity (FC), integrative and multimodal approaches to combine the two remain limited, especially for electrophysiological data. In the present work, we propose a new linear adaptive filter for estimating dynamic and directed FC using structural connectivity information as priors. We tested the filter in rat epicranial recordings and human event-related EEG data, using SC priors from a meta-analysis of tracer studies and diffusion tensor imaging metrics, respectively. Our results show that SC priors increase the resilience of FC estimates to noise perturbation while promoting sparser networks under biologically plausible constraints. The proposed filter provides intrinsic protection against SC-related false negatives, as well as robustness against false positives, representing a valuable new method for multimodal imaging and dynamic FC analysis.

## Introduction

The white matter architecture of the human brain constitutes the structural backbone for neuronal communication. A fixed network of axonal pathways wires an extremely rich repertoire of brain functions, from short-range interactions to large-scale dynamics that support perception, cognition, and action (Petersen & Sporns, 2015). As in all biological systems, properties of the structure constrain the possible functions. Organizational principles of structural brain networks, such as small-world and modular architectures, determine the topological space for functional interactions at the meso- and macro-scale (Hagmann et al., 2008; Sporns, 2010). At the microscale, the absence of a synaptic connection between two neurons makes a direct functional coupling biologically impossible. Despite the inherent link between structural (SC) and functional brain connectivity (FC), the two have been mostly investigated separately, and the potential benefits of integrative and multimodal approaches combining SC and FC remain largely unexplored (Lei et al., 2015).

The relationship between structural wiring and functional coupling is at the core of several statistical and biophysical models of brain networks (Honey et al., 2010). These models advocate a substantial overlap between SC and FC, both at the mesoscopic and macroscopic scales. Empirical and modelling studies on resting-state brain networks provide converging support, showing that the weights of structural and functional networks, as well as their topological features, tend to be correlated, and the strength of between-regions SC is typically a good predictor of their FC (Deco et al., 2013; Honey et al., 2009; Mišić et al., 2016; Skudlarski et al., 2008). Given the established overlap between structure and function, measures of FC may be meaningfully improved by taking SC into account, as suggested by a few proposed methods. In the framework of Bayesian modeling, for instance, structural graphs have been incorporated as priors for generative models of FC (Sokolov et al., 2019). A structural graph is typically obtained from in vivo diffusion-weighted imaging data (DWI) that quantify the anisotropy in the diffusion of water molecules along white matter tracts (Hagmann et al., 2008). The connectivity graph is either a binary or weighted undirected adjacency matrix that provides information about the presence and strength of a physical link between distinct brain regions. It has been shown how adding SC graphs as priors for effective connectivity substantially improves model evidence (Sokolov et al., 2019). Similarly, constraining FC for only present SC links and anatomically determined time lags may reduce false positives and improve the spatial resolution of electroencephalography (EEG) source imaging (Filatova et al., 2018; Takeda et al., 2019).

Whereas previous work has focused mainly on combining SC and FC for the analysis of functional Magnetic Resonance Imaging data (fMRI), similar integrative approaches are missing for the emerging field of time-varying directed FC analysis (Eichenbaum et al., 2021). Time-varying FC characterizes the dynamics of directed neuronal interactions that evolve at the millisecond scale, exploiting high-temporal resolution recordings, such as local field potentials and EEG source imaging data (Milde et al., 2010; Pascucci et al., 2018; Plomp et al., 2014). We here use the more general term FC througouth the paper, referring to these directed and dynamic measures of FC. Recently, we introduced a variant of the classic Kalman filter, the Self-Tuning Optimized Kalman filter (STOK; Pascucci et al., 2019), for modeling rapid changes in large-scale functional networks during evoked brain activity. Here, we present an extension of this algorithm that incorporates prior information on the structural connectivity: the structurally informed STOK (si-STOK). The algorithm provides a straightforward and novel tool to combine SC (e.g., DTI-derived metrics) with dynamic FC. We tested the algorithm in benchmark data and evaluated the effect of different SC matrices on the estimated FC. We demonstrated the advantages of combining SC with FC in terms of noise resilience and consistency of the estimates. We then compared the two algorithms in event-related large-scale functional brain networks during face processing. Our results showed that incorporating SC in dynamic FC promotes sparser and physiologically plausible topologies of functional networks, aiding the identification of the main network drivers and dynamics. Matlab and Python code for si-STOK are available on GitHub (https://github.com/PscDavid/dynet_toolbox; https://github.com/joanrue/pydynet).

## Results

### Somatosensory evoked potentials in rats

To incorporate structural priors in dynamic functional connectivity, we developed a variant of the self-tuning optimized Kalman filter (STOK) (Pascucci et al., 2019). The STOK is an adaptive filter that derives time-varying Multivariate Autoregressive coefficients (tv-MVAR) using a simple least-squares regression of present on past signals. We exploited this core feature of the filter to incorporate SC as shrinking priors of a least-squares solution (see Methods, Eq. [7]). SC matrices were used to calibrate the variance of a prior expectation of zero FC between each pair of nodes (Sokolov et al., 2019) (Figure 1). Strong SC values correspond to large prior variance, allowing FC to deviate from zero, whereas weak SC values reduce the prior variance and shrink FC toward zero. Hence, the filter’s estimates combine the strength of SC with the FC supported by the data (see Methods and Figure 1).

**Figure 1.**
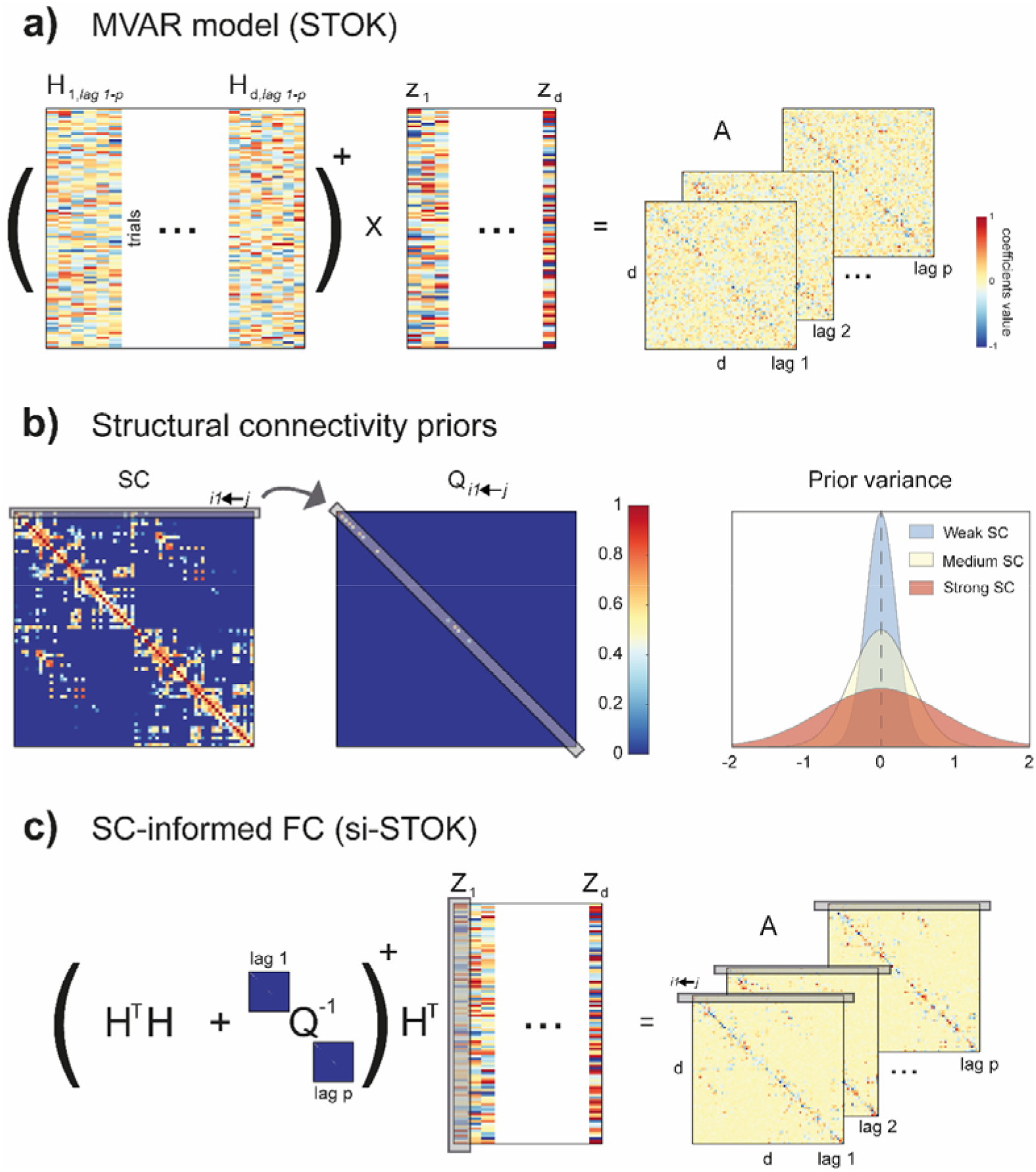
Estimation of structurally-informed dynamic FC using the si-STOK filter. a) A least-squares solution is used to estimate the coefficient matrix (A) of a time-varying Multivariate Autoregressive Model by regressing past (H) on present (Z) values of the multivariate time-series (see eq. [4], the symbol + stands for the matrix pseudoinverse). b) SC priors (e.g., DTI metrics) are incorporated in the filter as the prior variance on the expected zero FC from all the sender nodes to each receiver node. Weak SC corresponds to small prior variance, shrinking the coefficient estimates toward zero; strong SC corresponds to large prior variance, allowing the estimated coefficients to deviate more from zero when supported by the data. c) The regularizing matrix Q informs the least-squares solution with priors on the variance of autoregressive coefficients based on SC, resulting in MVAR models that combine FC and SC.

We tested the proposed algorithm, termed structurally informed STOK (si-STOK), on a benchmark dataset of epicranial EEG recordings in rats, from a whisker stimulation protocol (Plomp et al., 2014; Quairiaux et al., 2011). After whisker stimulation, action potentials originate and propagate rapidly (e.g., within 10-25 ms) from the contralateral primary sensory cortex, following the underlying structural connectivity (Plomp et al., 2014) (see Figure 2a). This pattern was accurately recovered by the STOK filter, which detected an overall larger Magnitude of outgoing Directed Influences (MDI, see Methods) from the contralateral sensory cortex (e4), at early post-stimulus latencies (13 ms, see Figure 2a and Pagnotta et al., 2018; Pagnotta & Plomp, 2018; Plomp et al., 2014). We used a directed, weighted matrix of SC that we derived from a meta-analysis of reported structural connections (Bota et al., 2015; Swanson et al., 2017) (see Methods). Compared to the regular STOK fileter, the inclusion of SC priors provided qualitatively similar results, with clearer dynamics and visible but minor changes (Figure 2a). The similarity between the results of the two algorithms was a consequence of the use of a dense SC matrix, with connection weights that did not deviate drastically from the estimated FC. When evaluating the magnitude of outgoing influences from e4 to all the other nodes, indeed, the results were highly comparable (Figure 2b). However, incorporating a sparser SC matrix, with only 25% of the strongest SC connections, led to evident changes. When one of the expected FC connections (from e4 to e2) was absent in SC, the resulting estimate decreased considerably (Figure 2b). Nevertheless, FC from e4 to e2 was still larger compared to two other SC-absent connections (from e4 to e1 and from e4 to e8) for which weak or no FC was supported by the data. Conversely, for connections with weak FC, strong SC did not drastically increase FC. This demonstrated that in the present modeling framework the inclusion of structural priors has a low risk of producing false negatives (the downscaling of FC for absent SC depends also on the strength of FC) while it is also robust against the risk of introducing false positives driven by strong SC in the absence of FC.

**Figure 2.**
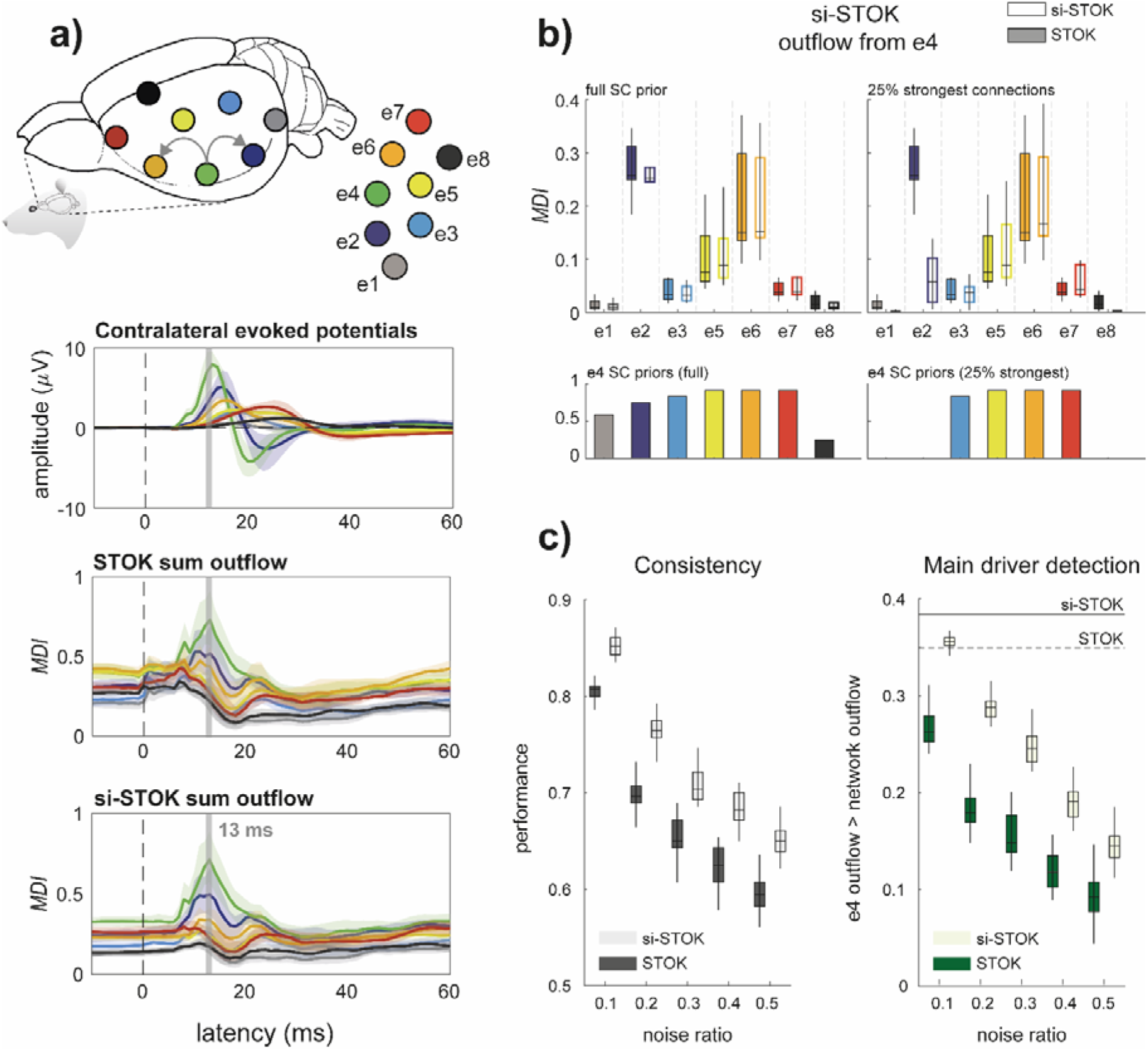
Validation in benchmark rat EEG data. a) Contralateral evoked potentials after whisker stimulation, showing the peak in the primary sensory cortex (e4). Activity propagates rapidly after stimulation from the primary sensory cortex to nearby parietal and frontal regions (e2 and e6). The role of the primary sensory cortex as the main driver of evoked activity is evident from the summed Magnitude of outgoing Directed Influences (MDI) estimated with both the STOK and the si-STOK filters. b) Effect of incorporating SC priors in the estimated directed influences from e4 to the rest of the network at the peak latency of the evoked activity (13 ms). Boxplots summarize the results of the STOK filter with (empty boxplots) and without (filled boxplots) SC priors, across ten animals. By using a dense SC matrix with almost uniform priors, the estimated FC are highly similar with and without SC priors. Retaining only 25% of the strongest SC connections demonstrates the relative shrinkage of SC-absent FC and the resistance of the algorithm against SC-related false positives: under an SC-absent prior, the expected FC from e4 to e2 was still larger compared to other connections, whereas strong SC priors did not inflate FC when weak FC was supported by the data (e.g., from e4 to e5). c) Performance evaluation under noise perturbations, after varying the proportion of signal to noise (additive) in the original data. Compared to the regular STOK, the si-STOK FC estimates showed overall larger consistency with the FC estimated in the absence of additional noise, as the proportion of noise perturbing the data increased (left panel). Similarly, the si-STOK showed a higher ability to detect the contralateral primary sensory cortex as the main driver of network activity at peak evoked latencies (right panel, the black line indicates the estimated e4 outflow, subtracted from the average network outflow at 13 ms, with the si-STOK in the absence of noise perturbations; the dashed line indicates the estimate obtained with the regular STOK). SC priors lead to an increased ability to detect e4 as the main network driver at all noise levels tested. Shaded lines in (a) are 95% CI of the mean.

To better appreciate the advantages of combining SC and FC, we compared the performance of the two algorithms under noise perturbations. We evaluated two criteria (see Methods): 1) the consistency of the estimated network at the e4 peak latency; 2) the ability to detect e4 as the main driver compared to the average network activity. The two criteria were tested by varying the ratio of noise to signal. As evident in Figure 2c, the si-STOK outperformed the regular STOK for both criteria at all the noise levels. This highlighted an additional important feature of the new filter: when FC is informed by SC, the estimated networks become more resilient to noise and present a consistent topology and nodal strength under perturbations.

### Human EEG data

After validating the algorithm in benchmark data recorded in rats, we employed the si-STOK filter to model FC in event-related human EEG data. We modeled FC in a large-scale network of 68 brain regions in response to faces and scrambled stimuli (Desikan et al., 2006) (see Methods and Figure 3a). On the electrode level, the comparison of evoked responses between faces and scrambled images revealed the typical topography and time-course of the face-related N170 component (faces minus scrambled, peak at 160 ms) with the largest difference between conditions localized in the right fusiform and nearby occipitotemporal areas (see Figure 3b).

**Figure 3.**
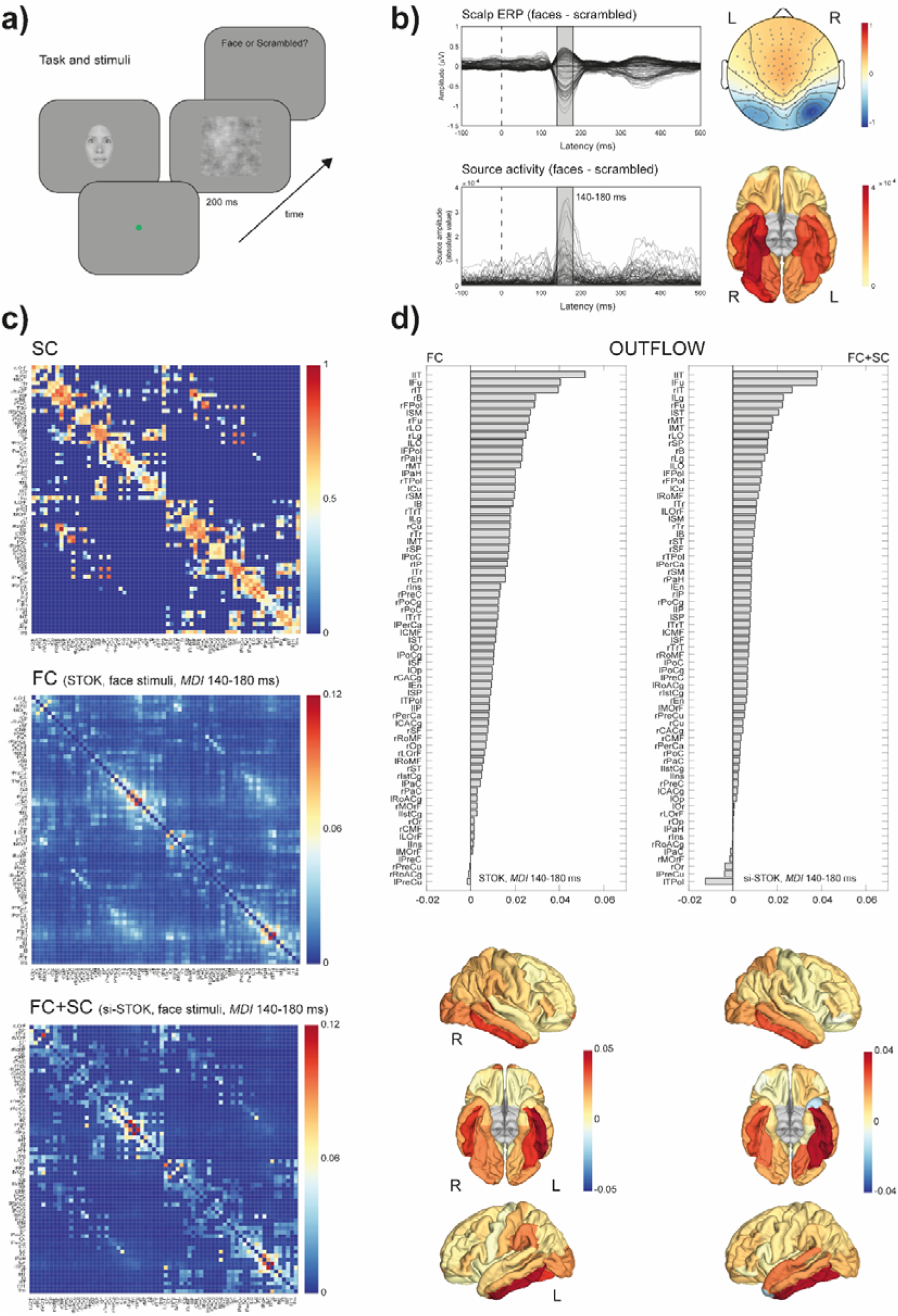
SC priors for large-scale FC analysis of human EEG data. a) Example of the sequence of events in one trial of the face detection task. Face and scrambled stimuli were randomly interleaved across trials. EEG data were time-locked to the onset of the stimuli. b) Scalp evoked responses showing the typical topography and time-course of the face-related N170 component (faces minus scrambled, peak at 160 ms, upper panels). Source reconstruction localized the source of face-selective responses in the right fusiform and nearby occipitotemporal areas (lower panels). c) The group SC prior matrix (upper panel) and the FC (MDI index) estimated with the regular STOK (central panel) and the si-STOK (lower panel) in response to face stimuli, averaged in a time window around the face-selective N170 response (from 140 to 180 ms post-stimulus). d) Ranked summed outflow in response to faces (faces minus scrambled) from all the 68 areas, obtained using the regular STOK (bar plot on the left) and the si-STOK (bar plot on the right). See supplementary File 2 for abbreviations. The inclusion of SC priors resulted in a less scattered topology of face-selective outflows, with the largest network drivers localized in primary, secondary visual areas and regions of the fusiform and inferior-temporal cortex.

Figure 3c shows the effect of incorporating SC priors on the estimated FC at a time window of interest around the N170 response. FC matrices obtained through the si-STOK filter showed the clear shrinkage of functional connections for weak and absent SC, leading to FC matrices that partly inherit the structure of SC but preserved intrinsic patterns of FC coupling (see Figure 3c), in line with the benchmark results. For the same time window of evoked activity, we compared the summed outflow from each area between the two conditions (faces minus scrambled, summed MDI) as a measure of changes in nodal strength during face processing. In the ranked outflow, the two filters agreed in identifying the bilateral inferior temporal gyrus and the left fusiform gyrus as the three areas with the largest increase in outflow in response to faces (see Figure 3d). Without structural priors, however, frontal regions were also ranked amongst the largest drivers at short post-stimulus latencies (e.g., rFPol, lFPol, see Supplementary File 2 for abbreviations) and the outflow increase in response to faces was more or less pronounced throughout the entire network. With the inclusion of structural priors, the ten largest drivers of face-related activity were all located in primary and secondary visual cortex, including the bilateral fusiform, lateral occipital cortex, lingual gyrus (e.g., V1), and regions in the temporal cortex (see Figure 3d), while the summed outflow from the rest of the network decreased progressively.

These comparisons suggest that the inclusion of SC priors refines the topology of FC networks and portrays the contribution of each node in a more physiologically plausible way. This feature can aid the identification of hubs and critical modules in large-scale FC analysis. A further important question is whether SC priors also affect the temporal dynamics of FC. We evaluated this aspect in a final analysis where we compared the estimated changes in directed influences in response to face and scrambled stimuli, time-locked to the stimulus onset (from -100 to 500 ms). For this analysis, we considered a subset of regions in the core face network: the right inferior temporal gyrus (rIT), the right fusiform (rFu), and the right superior temporal sulcus (rB). The functional role of these brain areas in face processing is well-documented (Fox et al., 2009; Haxby et al., 2000) and their SC is a predictive feature of face-specific activity (Saygin et al., 2011), representing a functionally specialized module with known structure-function relationships. Figure 4 shows the estimated time-varying MDI with and without SC priors. This comparison revealed clear differences that extended well beyond the basic outflow summary described above, specifically: 1) SC priors led to quantitative increases in unidirectional interactions at specific latencies, consistent with face-related evoked dynamics (e.g., from rFu to rIT, from rIT to rB); 2) SC priors more clearly show the sequence of unidirectional interactions between nodes (e.g., from rIT to rFu, followed by rFu to rIT); 3) SC priors shrunk and underestimated weak FC for SC-absent connections, reducing the risk of false positives (e.g., the connection from rIT to rB at very short post-stimulus latencies).

**Figure 4.**
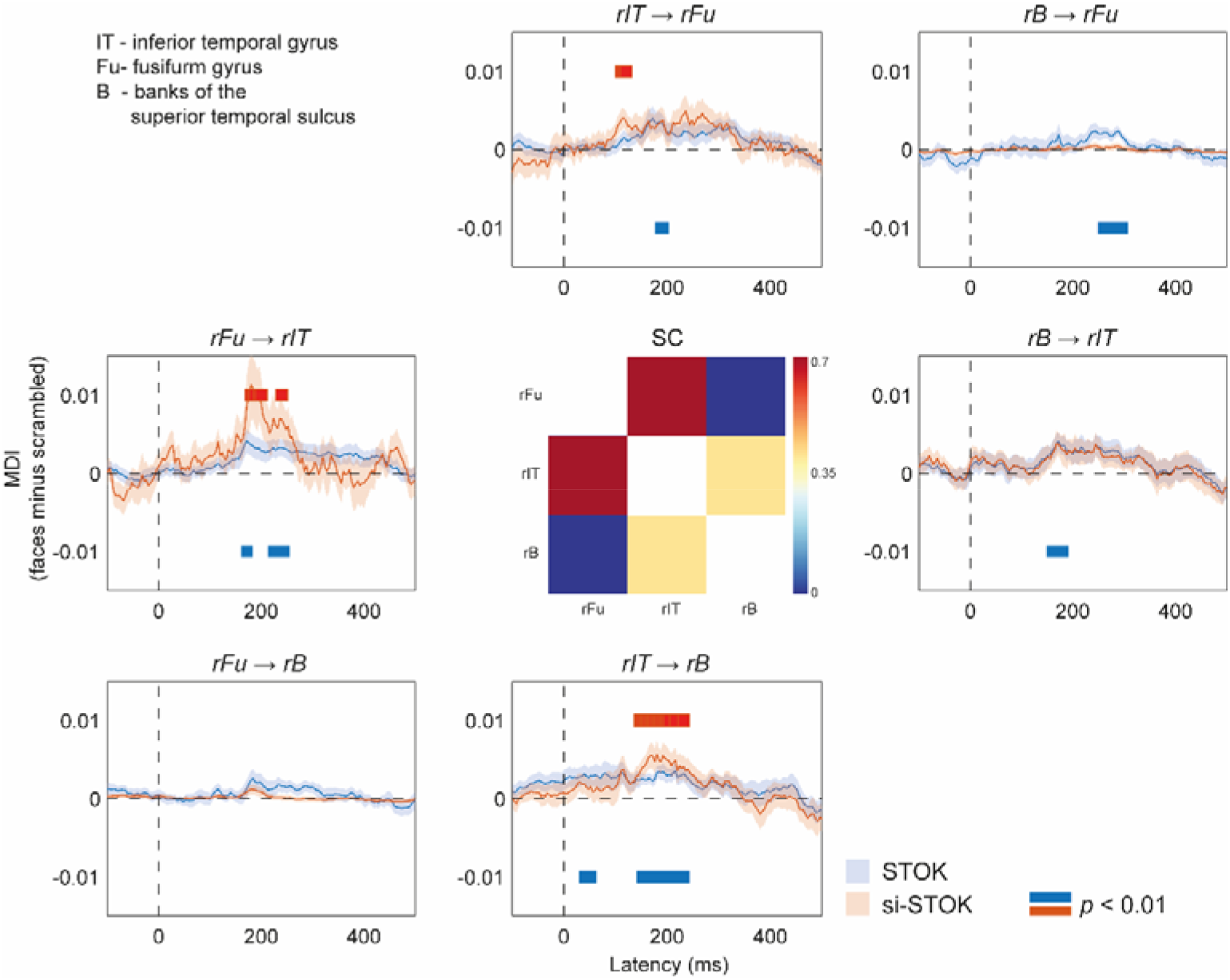
Effect of SC priors on the evoked dynamics of directed interactions in a subset of brain areas of the core face network. Shaded lines are standard errors of the mean. Red and blue horizontal bars highlights statistically significant differences (faces vs. scrambled, permutation test, *p*(unc.) < 0.01).

## Discussion

We introduced and validated a new adaptive filter for combining structural and dynamic functional connectivity, the si-STOK. The algorithm builds on a recent variant of the general linear Kalman filter (Kalman, 1960; Milde et al., 2010; Pascucci et al., 2019) and allows incorporating structural priors in a multivariate autoregressive modeling framework with high temporal resolution. We tested the filter in rat epicranial recordings and human EEG data, using SC priors from a meta-analysis of tracer studies and DTI metrics respectively. Compared to FC without SC priors, we found the following advantages. First, FC estimates were more consistent and more resilient to noise. Second, SC priors promoted sparser FC networks and favored a more accurate identification of the main network drivers at expected post-stimulus latencies. Third, using SC as prior variance provided solutions with intrinsic protection against SC-related false negatives (for discordant SC-FC, the filter relied more on the data and less on the prior), and robustness against false positives (strong SC did not inflate FC unless supported by the data).

The first two aspects represent desired features considering the expected sparsity of FC networks and the sparse topology of the underlying structural links (Markov et al., 2012; Pagnotta et al., 2019; Valdés-Sosa et al., 2005). Previous work has shown how sparse and regularized approaches to FC analysis can decrease spurious connections, increase robustness to noise (Pagnotta et al., 2019), and counteract issues due to limited data points (Antonacci et al., 2019; Valdés-Sosa et al., 2005). Informing sparse solutions through the fixed topology of SC links has the additional benefit of introducing a biologically grounded space for regularization. This represents an advantage in conditions where regularization may have a strong impact on the model structure, such as under multicollinearity and non-independence amongst time-series (e.g., under linear mixing of source EEG activity due to volume conduction; Anzolin et al., 2019; Haufe et al., 2013). The resulting FC partly inherits the sparsity and topological properties of the SC matrices, while preserving the strength and directedness of functional interactions. This may ultimately facilitate graph analysis of functional networks, such as the identification of FC hubs, modules, and nodal properties without additional sparsity-based and consensus-based thresholding, which may lead to unstable and threshold-dependent network estimates (Garrison et al., 2015).

The third important feature of the si-STOK is the protection against false positives due to invalid SC priors. Previous studies, including large-scale validations of tractography pipelines, have reported a high ratio of invalid connections and a substantial amount of false positives (Maier-Hein et al., 2017). This caveat undermines the possibility to simply mask or weight FC by SC, increasing the risk of inflating FC for invalid SC connections. The proposed algorithm employs the Generalized Tikhonov method (Plato & Vainikko, 1990), a powerful and versatile regularization scheme under a Bayesian perspective where priors are combined with the observed data. Strong SC, therefore, does not necessarily inflate FC, as demonstrated in our test in benchmark data. Conversely, strong FC can still be detected in the absence of SC. This was also evident from the results in rat EEG data, where we observed FC between the primary somatosensory and the parietal cortex (e4 to e2, see Figure 2) even under a strong prior of no SC. Although reduced, FC between these areas was still larger compared to other connections for which SC was present but only weak FC was expected from physiology (e.g., from e4 to e8). This accommodates the possible divergence between FC and SC connections (Honey et al., 2009; Lim et al., 2019), which may arise from indirect structural connections, false negatives in SC (Damoiseaux & Greicius, 2009), or because of the differential engagement of specific functional modules under different task demands (Sokolov et al., 2019).

From our test in benchmark data it is also clear that, under certain circumstances, SC priors have minor effects on FC. This may occur, for instance, when using dense SC matrices in conditions where the signal-to-noise ratio is high, or when SC does not deviate drastically from FC. The rat EEG data analyzed here are an example. The rat cortex is essentially flat (lissencephalic) with few expected deep sources. The signal recorded at each electrode is therefore an accurate representation of the activity flowing through the structural pathways in the cortex underneath (Quairiaux et al., 2011). Using dense SC priors, generally in agreement with the expected FC, provided no additional or diverging information (see Figure 2b). In many other applications, however, signals are contaminated with multiple sources of noise and the recordings are not direct measurements of brain activity and connectivity. Under these common circumstances, SC priors may have more appreciable and beneficial effects on FC estimates. Our manipulation of added noise confirmed this advantage (Fig 2c).

The proposed algorithm is the first to integrate SC in a directed and dynamic measure of FC. We evaluated the effect of SC priors on large-scale dynamics of directed interactions in human source EEG activity during a face detection task. We considered dynamic interactions in a subset of regions corresponding to key nodes of the core face network (Haxby et al., 2000; Saygin et al., 2011). Our results showed that SC priors can also shape the temporal dynamics of FC. Particularly, incorporating SC priors revealed an initial face-selective increase of FC from the right inferior temporal gyrus to the fusiform, followed by an increase of FC from the fusiform to the inferior temporal gyrus and from the inferior temporal gyrus to the superior temporal sulcus. Without SC priors, a precise sequence of FC changes was less distinguishable and face-evoked increases in FC were also present at unreasonably short post-stimulus latencies (e.g., from the inferior temporal gyrus to the superior temporal sulcus, see Figure 4). The observed pattern may well reflect the build-up of face-specific processing supported by a hierarchy of recurrent interactions in the core face network: the inferior temporal cortex may first relay information about global aspects (e.g., the shape and coarse structure of the face stimuli used here) to the fusiform which then feedback information about finer details (Goffaux et al., 2011; Sugase-Miyamoto et al., 2011; Tovée, 1995). Although the interpretation of these results remains speculative and goes beyond the purpose of the current work, the results of this analysis are a clear example of how SC priors can enhance or downregulate time-varying dynamics in FC.

In sum, we provide a new method for dynamic FC that incorporates priors on structural connectivity for the anaysis of multivariate electrophysiological signals. The algorithm offers a simple and powerful tool for multimodal imaging that can meaningfully contribute to integrative approaches in network neuroscience (Crimi et al., 2016; Lei et al., 2015; Sokolov et al., 2019; Stephan et al., 2009). It allows the incorporation of various types of SC, weighted or binary, symmetric or directed. Because of its simple form, the algorithm can flexibly incorporate different types of SC priors, from basic metrics, such as the number of white matter fibers and the Euclidean distance, to graph-derived metrics, such as the path length and communicability (Vázquez-Rodríguez et al., 2019), as well as FC priors from other modalities.

## Methods

### Benchmark EEG data

#### Rat EEG recordings

Benchmark data are publicly available EEG recordings from a grid of 16 stainless steel electrodes placed directly on the skull bone of 10 young Wistar rats (P21; half males). Data were collected during unilateral whisker stimulations under light isoflurane anesthesia (available from https://osf.io/fd5ru). Details about the recording can be found in the original publication (Plomp et al., 2014; Quairiaux et al., 2011). Data were acquired at 2000 Hz, bandpass filtered online, and down-sampled to 1000 Hz before connectivity analysis. All animal handling procedures were approved by the Office Vétérinaire Cantonal (Geneva, Switzerland) following Swiss Federal Laws.

#### Rat structural connectome

Structural priors for the rat EEG benchmark were obtained from a published meta-analysis of histologically defined axonal connections between cortical regions in rats (Bota et al., 2015; Swanson et al., 2017). Particularly, we used one dataset containing ranked connection weights based on reported association and commissural connections. Details of the dataset can be found in the original publication (Dataset S3; from Swanson et al., 2017). Ranked connection weights for pairs of regions corresponding to the electrodes recording sites were manually selected by an expert biologist from primary visual areas, somatosensory, primary, and secondary motor areas, and cingulate cortex (see Supplementary File 1). Connection weights, ranging from 0 to 12 (from absent to very strong, see Supplementary File 1) were organized into a 16-by-16 structural connection matrix whose main diagonal elements (e.g., self-connections) were set to the maximum value of 12. The structural matrix was then normalized to the maximum value and used as a prior for the time-varying connectivity analyses of the rat EEG data.

### Human EEG data

#### Task and stimuli

EEG data were recorded while twenty participants (3 males, mean age = 23 ± 3.5) performed a face detection task (see Figure 3a) in a dimly lit and electrically shielded room. Each trial lasted 1.2 s and started with a blank screen (500 ms). After the blank screen, one image (either a face or a scrambled image of a face) was presented for 200 ms and participants had the remaining 1000 ms to respond. The task was to report whether they saw a face or not (yes/no task) by pressing two buttons in a response box with their right hand (ResponsePixx, VPixx technologies). Faces and scrambled faces were randomly interleaved across trials. After the response and a random interval (from 600 to 900 ms), a new trial began. The experiment consisted of four blocks of 150 trials each, for a total of 600 trials, i.e., 300 with faces and 300 with scrambled faces. Face stimuli were female and male faces (4 by 4 degrees of visual angle, dva) taken from online repositories and cropped with a Gaussian kernel to smooth the borders. Scrambled images were obtained by fully randomizing the phase spectra of the original images (Ales et al., 2012). Stimuli were generated using Psychopy (Peirce, 2008) and presented on a VIEWPixx/3D display system (1920 ×□1080 pixels, refresh rate of 100□Hz). All participants provided written informed consent before the experiment and had a normal or corrected-to-normal vision. The experiment was approved by the local ethical committee.

#### EEG acquisition and preprocessing

Data were recorded at 2048 Hz with a 128-channel Biosemi Active Two EEG system (Biosemi, Amsterdam, The Netherlands). Signal quality was ensured by monitoring and maintaining the offset between the active electrodes and the Common Mode Sense - Driven Right Leg (CMS-DRL) feedback loop under a standard value of ±20 mV. After each recording session, individual 3D electrode positions were digitized using an ultrasound motion capture system (Zebris Medical GmbH). One participant was excluded due to too many motion artifacts, leaving 19 datasets for analysis. Further details on the recordings and preprocessing pipeline can be found in the original manuscript of the VEPCON dataset (OpenNeuro Dataset ds003505; Pascucci et al., 2021).

#### EEG source imaging

EEG source imaging was performed using Cartool (Brunet et al., 2011) and custom-made scripts in Matlab R2020b (9.9.0.1524771 Update 2). Source reconstruction was based on individual MRI data and the LAURA algorithm implemented in Cartool (regularization 6; spherical model with anatomical constraints, LSMAC), limiting the solution space to grey matter voxels. Source activity for freely oriented dipoles was extracted from all the source points inside each of the 68 cortical areas and projected to a representative single direction for each area, using the singular values decomposition approach (Rubega et al., 2018; time window for estimating the main direction: 140-250 ms post-stimulus). Before functional connectivity analysis, a global z-score transformation was applied to the entire dataset of each participant. Epochs of source activity corresponding to trials with behavioral errors were then removed and the dataset was divided into two conditions, according to trials containing faces or scrambled stimuli.

#### MRI acquisition and preprocessing

A detailed description of the MR acquisition and preprocessing can be found in the original manuscript of the VEPCON dataset (OpenNeuro Dataset ds003505; Pascucci et al., 2021).

#### Structural connectome

Structural connectivity matrices were estimated from the reconstructed fiber orientation distribution (FOD) image using the SD_stream deterministic streamline tractography algorithm implemented in MRtrix 3.0.0-RC1 (Tournier et al., 2019). Fiber streamline reconstruction started from seeds in the white matter that were spatially random, and the whole process completed when 1M fiber streamlines were reconstructed. At each streamline step of 0.5 mm, the local FOD was sampled, and from the current streamline tangent orientation, the orientation of the nearest FOD amplitude peak was estimated via a Newton optimization on the sphere. Fibers were stopped if a change in direction was greater than 45 degrees. Fibers with a length not in the 5-200 mm range were discarded. The streamline reconstruction process was complete when both ends of the fiber left the white matter mask. Then, for each scale, the parcellation was projected to the native DTI space after symmetric diffeomorphic co-registration between the T1w scan and the diffusion-free B0 using ANTs 2.2.0. Finally, the connectivity matrix was built according to the Desikan parcellation atlas, using the log of the number of fibers as the connectivity measure. Only 68 areas from cortical volumes were included in the structural matrix and used for functional connectivity analysis. A consensus group-representative structural brain connectivity matrix was generated from the connectomes of all participants connectomes using the method introduced in (Betzel et al., 2019). Each particpiants’ connectivity matrix was then thresholded by preserving the group-representative connection density independently for intra- and interhemispheric connections. This allows retaining more inter-hemispheric connections in comparison to simple connectome thresholding. The resulting connection density is set to 30%. The median of the obtained structural connectomes across participants was then normalized to its maximum and used as a group structural prior for connectivity analysis.

### Adaptive filtering

#### Self-Tuning Optimized Kalman filter (STOK)

To incorporate structural priors in dynamic functional connectivity we used a linear adaptive filter, the Self-Tuning Optimized Kalman filter (STOK; Pascucci et al., 2019), as the base algorithm. STOK is a high-temporal resolution and noise resilient filter for modeling time-varying multivariate autoregressive processes (tv-MVAR) of the form:

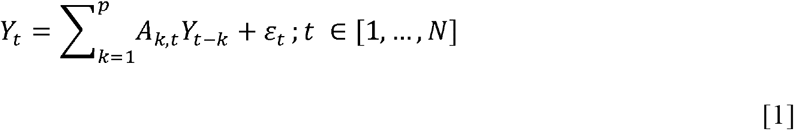

where *Y* is a multi-trial and multivariate set of *d* time series of dimension [trials x *d*] (e.g., activity signals from different brain regions),*t* refers to time samples (with *N* the total length of the time segment considered), *A*_*k,t*_ are [*d x d x P x N*] matrices of autoregressive coefficients for each lag *k* of a chosen model order *P, ε*_*t*_, is zero-mean white noise with covariance matrix *∑*_*ε*_ (also called the *innovation* process). Eq. [1] can be represented in the following state-space form:

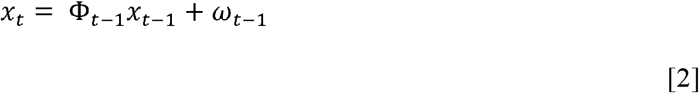

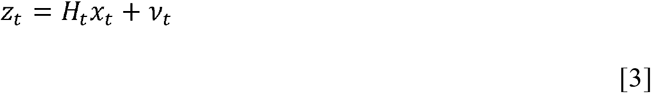

where Eq. [2] represents the latent state *x*_*t*_(e.g., the MVAR process) as a random walk from the previous state *x*_*t*−1_ [*d x d x P*], with transition matrix *Φ*_*t*−1_ and uncorrelated zero-mean noise *ω*. In Eq. [3], the observed data *z*_*t*_, are expressed as a linear combination of the latent state *x*_*t*_ and a projection matrix *H*_*t*_, under white noise perturbation *v*_*t*_. The link with Eq. [1] is established by recursively defining *H*_*t*_ from the past of the time-series in *Y* (from, *t-1* to the model order *P*), and *z*_*t*_, as the values of *Y* at present time, *t*. This leads to the least-squares estimate:

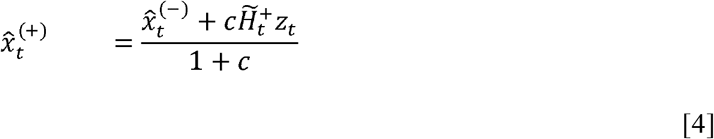

in which the recursive update of 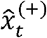 is a weighted average of the previous state 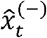 and a least-squares reconstruction from recent measurements 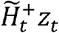. The matrix 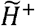 is the damped Moore-Penrose pseudoinverse (^+^) of *H*, in which small singular values are attenuated to retain a prespecified portion of the variance (here we retained 0.99 of the variance for the rat EEG data and 0.9 for the human EEG data). The variable *c* is a self-tuning adaptation constant that automatically updates the speed of the filter depending on its residuals. The complete derivation of the STOK can be found in (Pascucci et al., 2019).

#### Structural priors

The prior information is incorporated in the filter as a regularizing operator. Recall that the update of 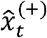 in Eq. [4] —i.e., the estimated matrix of tv-MVAR coefficients, requires the ordinary least-squares solution to 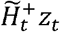:

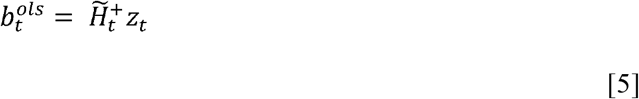

with 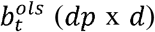 representing the matrix of autoregressive coefficients that updates the previous state 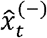. A principled way to incorporate priors in Eq. [5] is the use of a *Generalized Tikhonov regularization*, which admits the closed-form solution:

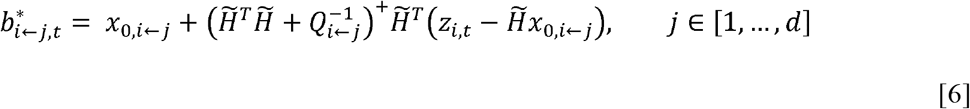

Where *x*_0_ is the expected value of *b*, and *Q*^− 1^is the inverse covariance matrix, or *precision matrix*, of *x*_0_. 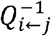 acts as a regularizing or Tikhonov matrix in Eq. [6]. The inclusion of a regularizing matrix allows specifying penalties on the estimated coefficients. When the regularizing matrix is a multiple of the identity matrix, the solution corresponds to the classical *L*_2_ norm. When the main diagonal contains distinct elements, however, the solution penalizes the coefficients differently, depending on the strength of the corresponding value in the regularizing matrix. This form of regularization offers a straightforward solution for incorporating structural priors in a tv-MVAR model. By solving Eq. [6] for each channel separately (e.g., for each signal in the multivariate time series), elements on the main diagonal of *Q* ^*−1*^ can be used to penalize the inflow from channel *j* to channel *i* (*i←j*) depending on structural priors, that is, an a priori structure can be imposed on the contribution of all the channels to the activity observed in each one.

Although this approach is deterministic, it has a natural Bayesian interpretation: Eq. [6] is equivalent to expressing a prior belief on the functional connections *b*_*i←j*_ entering each channel, under a multivariate normal distribution *N*(*b*_*i←j*_*;x* _0_,*Q*_*i←j*_). This regularization scheme requires the non-trivial conversion of SC to FC priors. A commonly employed strategy is to set the prior expectations on *x*_0_ to zero (e.g., no functional connectivity, Sokolov et al., 2019; Stephan et al., 2009), and to define the prior variance *Q*_*i←j*_ based on the strength of SC. Under the mild assumption of a positive and monotonic relationship between structure and function, strong SC can be translated into large FC prior variance 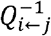, corresponding to small regularization values in 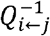 that favor non-zero FC when supported by the data. Conversely, weak SC can be translated into small prior variance, increasing the effect of regularization, and shrinking the tv-MVAR coefficients toward their expected value of zero. Mapping SC to FC priors also requires the scaling of SC to a range of suitable values, given that regularization acts on the magnitude and scale of autoregressive coefficients. Normalized SC values are in the 0-1 range. For the sake of the present work, we scaled SC from 10^−4^ to 0.1, a range that produced clear effects of regularization for both rat and human EEG data, without an excessive shrinking of all the coefficients.

Hence, the Tikhonov matrix 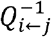 in Eq. [6] is a diagonal matrix whose non-zero elements represent the inverse prior variance for the functional connectivity between *j* input sources and a receiver node *i*. By setting *x*_0_ equal to zero, we obtain:

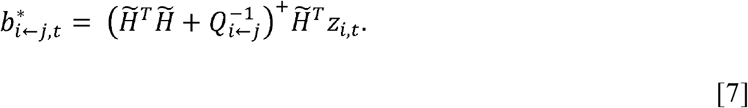

In a Bayesian view, the precision matrix 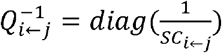 in Eq. [7] determines the extent to which the posterior estimate (e.g., the functional connectivity between two nodes) can deviate from its expected value of zero. As a result, the estimated matrices of tv-MVAR coefficients combine information from both functional and structural connectivity: weak SC decreases the prior variance and increases regularization (e.g., expressing a strong belief that a functional connection is likely to be absent, to reduce false-positive connections); strong SC increases the prior variance and decreases regularization, allowing functional connections to deviate from zero when supported by the functional data.

The new coefficients 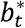 obtained from Eq. [7] (posteriors) are then substituted in Eq. [4], leading to the recursive update equation with structural priors:

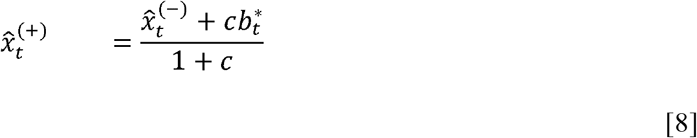

which ultimately provides an estimate of tv-MVAR coefficients *A* informed by the properties of SC. The structural connectivity matrix can be either symmetric or asymmetric (e.g., for unidirectional priors), binary or weighted, with values that increase as a function of the expected strength of a connection between two nodes.

### Time-varying directed functional connectivity

#### Directed connectivity measure

In the analysis of FC, we used a measure of directed connectivity derived from the estimated matrices of tv-MVAR coefficients. This measure, that we termed Magnitude of Directed Influence (MDI), corresponds to the magnitude of autoregressive coefficients over lags, and quantifies the time-domain unidirectional influence from a sender *j* to a target node *i*:

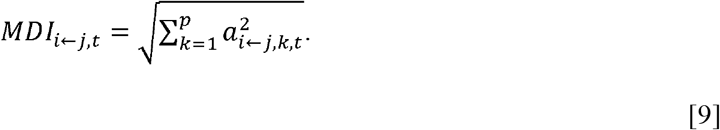

A similar measure, called *direct causality* (DC), has previously been used to estimate the amount of direct causal influences in multivariate systems (Kamiński et al., 2001; Porcaro et al., 2009).

#### Rat EEG functional connectivity

We tested the new algorithm and compared the performance against the regular STOK using benchmark EEG data from epicranial recordings in rats. These data provide a good benchmark for time-varying connectivity analysis because of the known structural and functional connectivity patterns. After whisker stimulation, the evoked FC is expected to follow the underlying SC, with the activity that propagates from the contralateral primary somatosensory cortex to nearby parietal and frontal regions, at short latencies (5-25 ms post-stimulation). This pattern was extensively validated in previous work (Pagnotta & Plomp, 2018; Pascucci et al., 2019; Plomp et al., 2014).

As a proof of concept, we first verified the effect of adding structural priors with different levels of sparsity. Two SC matrices were used as priors for the si-STOK, one obtained from the original SC (see Rat structural connectome and Supplementary File 1), the other obtained by applying proportional thresholding to the same SC, retaining only 25% of the strongest connections, setting the remaining to zero and the diagonal (self-self connections) to one. The *MDI* metric was derived from tv-MVAR models with a model order of 4 (Pagnotta & Plomp, 2018; Pascucci et al., 2019). In the comparison, we focused on the outflow from the contralateral somatosensory cortex to the rest of the network, at the peak latency. The peak latency was estimated from the results of the regular STOK as 13 ms post-stimulus, in line with previous reports (Pascucci et al., 2019; Plomp et al., 2014).

In a second step, we compared the performance of the two filters under noise perturbation. We mixed the original data with white noise signals of the same size as the data and with an amplitude corresponding to the 95^th^ percentile of the data. We varied the mixing ratio between the original data and the noise signals in five levels (from 0 to 0.5, in steps of 0.1; 0 = 100% of the data and 0% of the noise). The noise at each level was regenerated 30 times. For each iteration, we estimated: 1) the difference between the outflow from the contralateral somatosensory cortex and the average outflow of the network, at peak latency, and 2) a binary, directed adjacency matrix preserving 50% of the strongest connections at peak latency. From the first measure, we estimated the difference between the outflow from the primary somatosensory cortex and the network average outflow, across animals —i.e., the ability of the two algorithms to discriminate the somatosensory cortex as the main driver under noise perturbations. From the second measure, we derived a metric of consistency of the estimated networks under increasing noise. The consistency was obtained as the proportion of binary connections in the adjacency matrix that, for each level of noise larger than 0, were identical to those estimated in the absence of noise —i.e., the consistency of the estimated strongest connections under increasing noise. For both measures, the zero-noise level was used as a baseline.

#### Human EEG functional connectivity

In the analysis of human EEG data, we evaluated the results of the STOK and si-STOK filter in estimating large-scale FC during face processing (see Figure 3a). For both filters, we used a model order of ten, in line with values used before (Pascucci et al., 2018, 2019). We first compared evoked activity after face and scrambled stimuli at the scalp and source level (see Figure 3b). We selected a time window centered on the scalp and the source N170 peak component (140-180 ms) as the latency of interest for FC results. In the N170 window, we evaluated the estimated difference in network MDI across conditions. For this difference, we compared the summed outflow obtained with the two filters, from each node of the 68-areas network (see Figure 3d). In a subsequent analysis, we focused on dynamic directed interactions among a subset of areas known to be involved in face processing (e.g., regions of the core face network, Haxby et al., 2000). We considered three areas, all in the right hemisphere: the fusiform gyrus (rFu), the inferior temporal gyrus (rIT), and the superior temporal sulcus (rB) (see Figure 4). We then compared directed influences among each pair of areas in response to face or scrambled stimuli across time. Significance was assessed using group-permutation statistics, as the proportion of group-averaged MDI differences that were larger or smaller than the observed ones, after shuffling the sign of the difference across participants 100000 times.

## Acknowledgements

The authors thank Guru Prasad Padmasola and Charles Quairiaux for providing guidelines and useful material for the analysis of the rat benchmark data, and Larry W. Swanson and Joel Hahn for providing the rat structural connectivity dataset.

